# The promise of AlphaFold for gene structure annotation

**DOI:** 10.1101/2025.10.21.683479

**Authors:** Helen Rebecca Davison, Ulrike Böhme, Shahram Mesdaghi, Paul A. Wilkinson, David S. Roos, Andrew R. Jones, Daniel J. Rigden

**Affiliations:** Department of Biochemistry, Cell and Systems Biology, Institute of Systems, Molecular and Integrative Biology, University of Liverpool, United Kingdom; Computational Biology Facility, University of Liverpool, United Kingdom; Department of Biology, University of Pennsylvania, Philadelphia, PA 19104, USA

**Keywords:** bioinformatics, proteomics, structural prediction, quality metrics, annotation

## Abstract

**Background:** As sequencing technology improves, more genomes become available. Most lack annotation, automated methods are error prone, and few genomes are ever manually curated due to time and cost. Protein structure prediction software may provide new angles for assessing and improving gene models without requiring experimental data. In this paper, we explore whether scores from protein structure prediction can aid in scoring gene model quality.

We chose three species (*Fusarium graminearum*, *Toxoplasma gondii*, and *Aspergillus fumigatus*) from the VEuPathDB database which have collectively undergone more than 1000 manual curation events. We modelled translations of the gene models with AlphaFold 3, before and after curation, collecting various scores. Then we carried out structure searching of the PDB with Foldseek and sequence-based domain identification using InterProScan. We profiled the scores produced by these methods to identify those best for gene model assessment.

**Results:** AlphaFold 3 scores strongly favoured manually improved over pre-improvement models, supporting 75% of manually-curated changes in *F. graminearum*, 65% in *T. gondii*, and 84% in *A. fumigatus* (the lower percentage in *T. gondii* attributed to a high level of disorder). Further, combining scores across multiple tools (AlphaFold 3, Foldseek and InterProScan) provided additional improvements in model scoring.

**Conclusion:** Overall, the most discriminative scores combined outputs of AlphaFold 3 and Foldseek. Our results therefore highlight the potential of scores derived from deep learning-based protein structure prediction for scoring gene models in the absence of experimental data. Future work should focus on intrinsically disordered regions and developing integrated tools to apply this approach.

## Background

The growing accessibility of genome sequencing means that genome assemblies are increasingly available for many organisms and isolates yet gene structure annotation remains a challenge. Since the beginning of genome sequencing efforts, extensive manual curation of model organisms has corrected exon-intron structure and start codons genome-wide (1). However, the efficacy of manual curation is dependent on the availability of large amounts of high quality experimental data and highly trained and specialised curators. Many of the same challenges for accurate next-generation gene annotation persist after decades of work (2). Newly sequenced genomes for less well studied organisms rarely have the depth of experimental data, nor funding for curators to take a gene-by-gene approach to improvements (3). Thus, alternative approaches will be required for the increasing number of species whose genomes have been sequenced but for which no functional genomics datasets are available (4) or Darwin Tree of Life (5).

Genome-wide, automated eukaryotic gene finding software were first developed in the mid 1990s, including GeneMark (6,7), GENSCAN (8) and later AUGUSTUS (9), using intrinsic genomic signals for the identification of coding regions These packages were later incorporated into - or superseded by - methods using signals from protein sequences (*e.g.*. orthologs), or experimental data (initially ESTs, and later RNASeq *e.g.* MAKER (10), BRAKER1/2/3 (10,11), Gnomon/NCBI (12), Ensembl pipelines (13) and more recently, deep learning pipelines such as Helixer (14)). All pipeline methods that incorporate multiple types of evidence have some internal methods for ranking plausible transcript models. For example, BRAKER2/3 uses TSEBRA to combine evidence from protein sequence orthology and RNASeq, thereby achieving better performance than with a single evidence type (11,15). BRAKER3 is considered state-of-the-art capable of reaching F1 scores of 50-80%, but still yields a high number of predictions that may not reflect biological reality (11,16). A few pipelines produce user-accessible scores that can be used to evaluate the trustworthiness of gene models. MAKER2 and FINDER provide a per-model quality score called Average Edit Distance (AED) (17,18). AED is determined by the distance between a given gene model and the assembled transcript evidence from RNASeq. Other methods have been developed to allow gene model scoring independent of gene finding software. For example, TRaCE uses AED, InterPro domain coverage and protein length, applied in a simple voting system (19). PSAURON uses a neural network trained on NCBI RefSeq genomes to score the plausibility of gene models (20).

Here we propose that the latest generation of deep learning-based protein structure prediction methods could provide fresh and complementary sources of information to support gene model prediction. The difficulties inherent in accurately identifying exons can result in missing or supernumerary exons which impact the integrity and plausibility of the 3D protein structure in ways we suggest should be detectable in protein models. Programs like AlphaFold 3 have revolutionised protein structural bioinformatics by enabling fast, accurate modelling of most proteins (21–23). These tools, and other similar methods, provide reliable estimates of confidence in structural predictions, which is key to their utility and broad adoption.

AlphaFold 3, which is our focus in this paper, produces three main scores: predicted local distance difference test (pLDDT), Predicted Aligned Error (PAE), and predicted Template Modelling score (pTM). The pLDDT provides a per residue estimate of the probability that the local environment of a given amino-acid in the model resembles that of the same position in the true structure. Regions with low pLDDT scores can result not only from unreliable prediction, but can also authentically reflect intrinsically disordered regions of the target protein that lack stable structure, at least in the absence of binding partners (24). The PAE provides confidence in residue-residue separations for the whole structure, estimating the likely error on all residue pairs. And finally, the pTM is intended to provide an overall score for fold similarity (on a scale from 0 to 1, where 1 means identical structure) between the model and the true structure. One important note is that the pTM is depressed by the presence of low-confidence (*e.g.* intrinsic disorder) regions in the model (25). Our preliminary results using AlphaFold 3 predictions from alternative gene annotations in the Asian rice (*Oryza sativa*) genome suggested that this approach is worth exploring (Figure 1 shows an example). They demonstrate that subtle incorrect splicing signals can be detected in AlphaFold structures.

**Figure 1.**
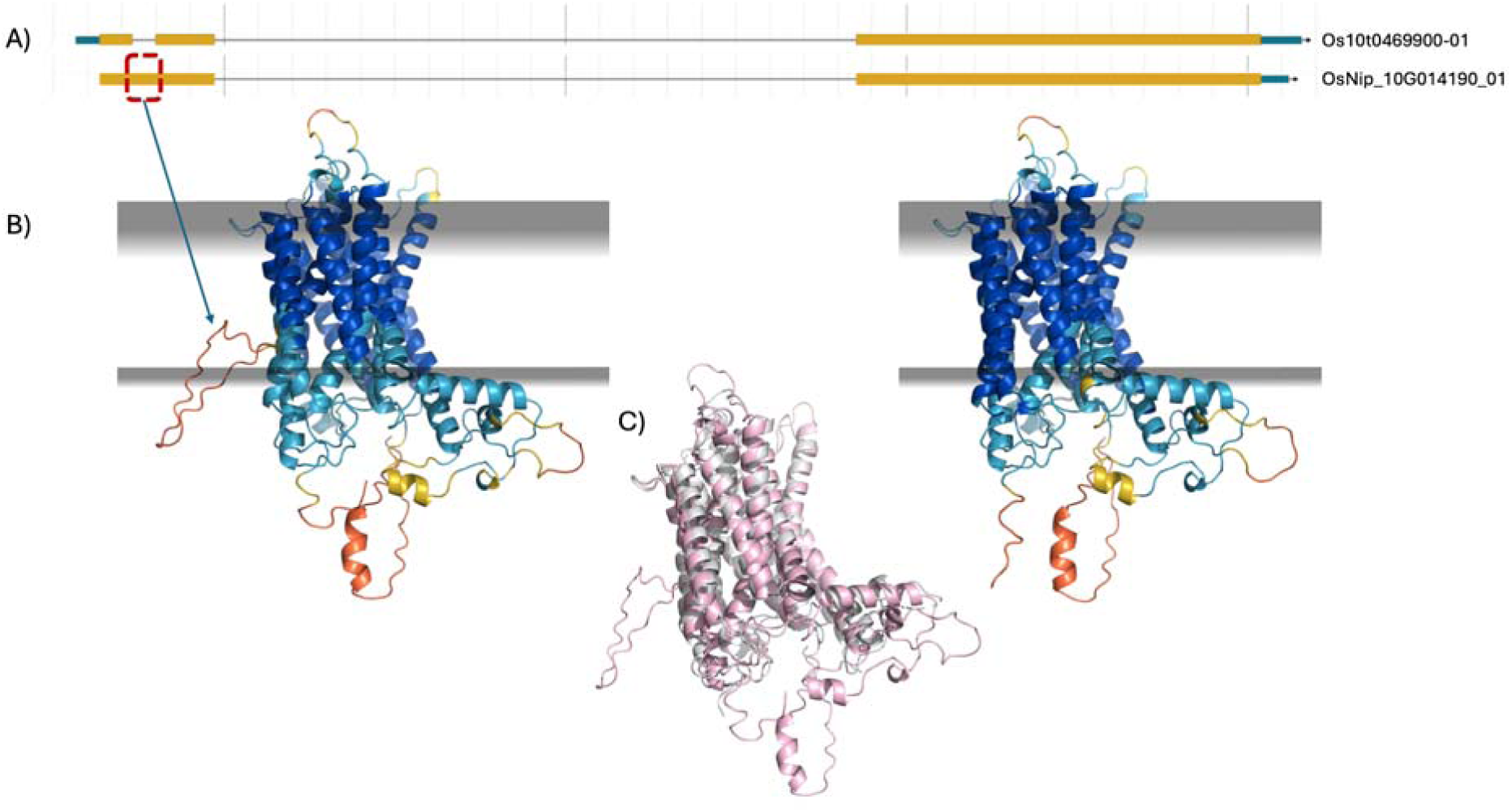
An illustrated example of how structure helps assess gene model quality. *a)* Two gene models for Asian rice (Oryza sativa) TGF-beta receptor, type I/II extracellular region family protein from different annotation sources. OsNip_10G014190_01 (new gene model generated with long-read RNASeq from Gramene.org) predicts two exons, Os10t0469900-01 (RAP-DB) predicts three exons. b) AlphaFold2 (ColabFold) structural models of OsNip_10G014190_01 and Os10t0469900-01. The models highlight a putative intron read-through that generates a low-confidence loop inconsistent with the tightly bundled α-helical structure. Grey planes represent predicted membrane boundaries calculated by the PPM server (26). Notably, the introduced loop is positioned within the membrane, providing a clear indication of misannotation. Model confidence is indicated by pLDDT colouring, ranging from blue (high confidence) to red (low confidence). c) Structural alignment of the OsNip_10G014190_01 model (pink) with its closest structural neighbour identified by Foldseek, PDB entry 4oh3 (grey). The model extends through an intron, generating an erroneous loop absent from the structural database hit.

We do not expect that scores from AlphaFold 3 alone will always unambiguously identify the correct version of a gene amongst alternatives. Where a structure prediction strongly resembles an experimentally characterised protein it can be considered more likely to result from a correct gene model. Therefore, we further propose screening AlphaFold 3 results against databases such as the Protein Data Bank with Foldseek or InterPro with InterProScan (27–30) (Figure 1).

In this paper, we test this novel application of AlphaFold 3 against the results of recent manual gene model curation exercises in three organisms (*Fusarium graminearum* str. PH-1, *Toxoplasma gondii* str. ME49, and *Aspergillus fumigatus* str. Af293). The organisms are held in the VEuPathDB database (31), and each can be considered a challenge for gene annotation. *Toxoplasma*, while well studied and -annotated, is known for its highly disordered proteins, typical of *Apicomplexa* (32). Fungi in general have hugely varying and complex genome structures, and tend to have higher gene density and fewer introns compared to other eukaryotes (33). All three species were chosen because they have undergone substantial manual curation and have metadata records detailing the rationale in each case. We compare scores from AlphaFold 3, Foldseek, and InterProScan and find that each has excellent discriminatory power across all three organisms. Importantly, we find that there is complementarity between the scores indicating that their combination with existing metrics in a machine learning framework could be useful for improving the performance of the next generation of gene finding tools.

## Results

### Overview of manually curated gene structure annotations

Our dataset comprised 444, 317 and 461 gene pairs, respectively from *Toxoplasma Gondii str. ME49, Fusarium Graminearum str. PH-1* and *Aspergillus fumigatus str. Af293,* listed in Supplementary table 1. We filtered the full gene models sets for pairs with <80% sequence identity to exclude minor changes in coding sequence. Old gene models are a mix of *in silico* and historical manual annotations. New models were exclusively manually curated over a number of months by Böhme.

The manual curation resulted in various kinds of changes to the structural annotation (Supplementary data 1-3). In extreme cases, a gene model was deleted, or was created for a region not previously annotated. In other cases, two gene models were combined or conversely, a single model split into two distinct coding units. However, in most cases the change resulted in the inclusion or elimination of one or a small number of exons, a category we call ‘Changed’ - and the focus of this analysis. This category is where we expected a protein structural perspective to be most useful: scores would detect the incompleteness of a structural domain, or the distortion and packing defects resulting from improper incorporation of a translated intron ( Figure 1). Comparing new versus old annotations showed that there was a slight tendency towards longer models post-improvement (Supplementary figure 1a), with variable patterns in the numbers of Coding DNA Sequences (CDSs, Supplementary figure 1c).

The extent of intrinsic disorder in our three focal species varies considerably, as assessed by Metapredict3 (Supplementary figure 1b). For example, the median (of the percentage disorder across each amino acid per protein) across all proteins is ∼20-30% in the new annotations of the two fungi, but much higher at ∼50% in *T. gondii*. Interestingly, the distributions of disordered content before and after manual curation vary considerably (Supplementary figure 1b) suggesting that many re-annotations impact disordered regions, potentially challenging our structure-based method.

### Signals from structural prediction

We explored a variety of scores reflecting structural quality in the AF3 models (see Methods). Median pLDDT, pTM and mean PAE all preferred the new gene annotations over the old versions (Figure 2, 3 and 4, Supplementary figure 2). Additionally, we found that counting the number of confidently modelled residues (pLDDT greater than a threshold) provided the clearest signal (Figure 2, 3 and 4, Supplementary figure 2).

**Figure 2.**
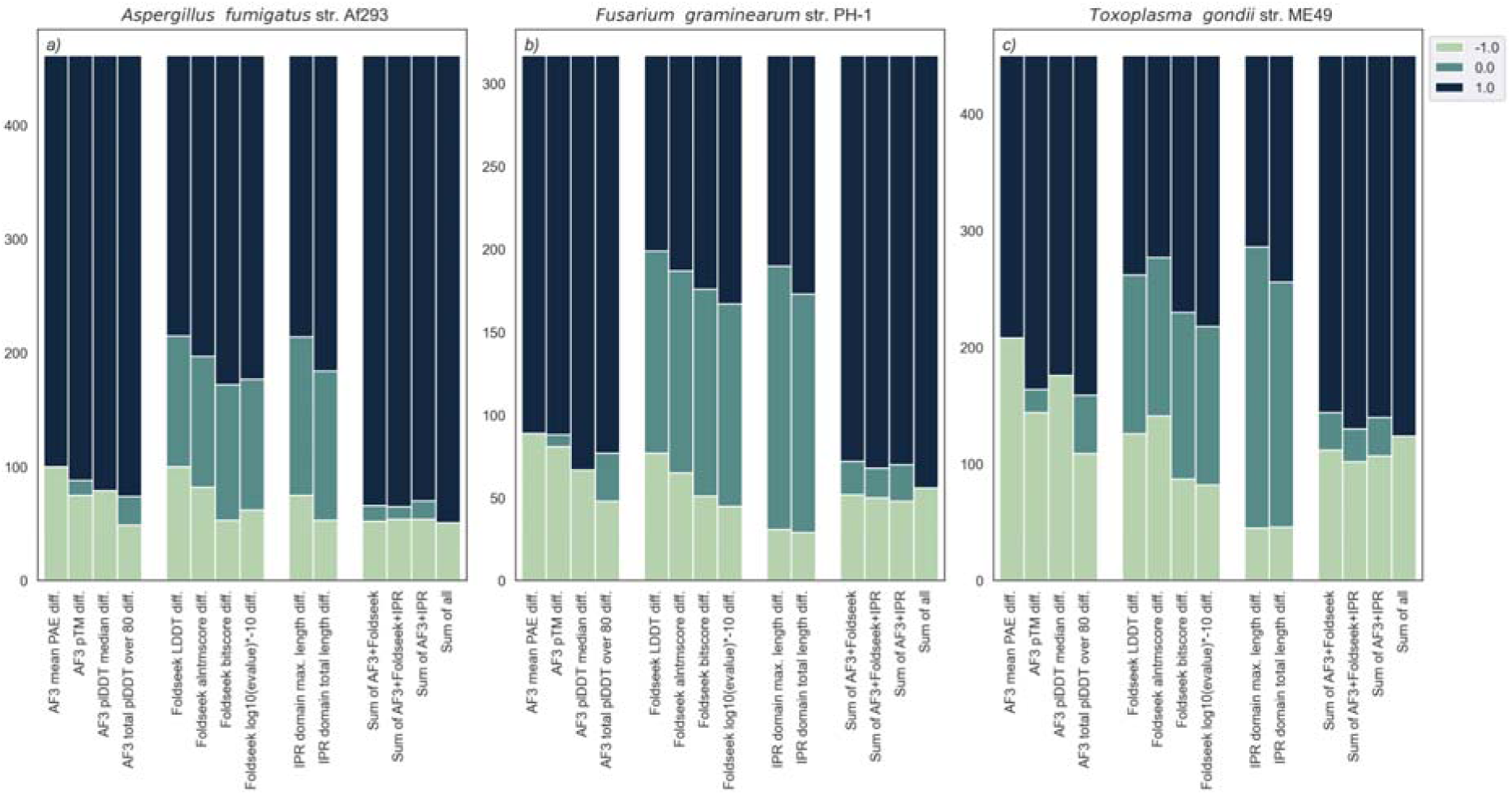
Stacked bars of difference values for each score grouped by source (AF3, Foldseek, IPR) Positive changes i.e. where the score favours the new gene model, are assigned +1, no change is 0, and negative change is −1. The rightmost group of bars in a), *b)* and c) show the summed MaxAbs transformed difference scores of combinations of the best AF3 (number of residues with pLDDT >80), Foldseek (Bitscore), and/or InterProScan (IPR total length) metrics.

In particular, counts of residues that score above 80 pLDDT and above 90 appear to be highly discriminative (Figure 3 and 4, Supplementary figure 2). The poor performance of pTM can be explained by its correlation with intrinsic disorder (Supplementary figure 3.). This aligns with the observation that the pTM of the same structured domain can be depressed by the simple addition of disordered termini (25). Similarly, the fact that median pLDDT is less discriminatory than the count of high pLDDT residues (Figure 1 and 4) can be explained by the fact that the median will be reduced by the presence of disorder outside of any folded domains, whereas the count will not. We expect structure modelling to offer clear discrimination principally in the folded structure.

**Figure 3.**
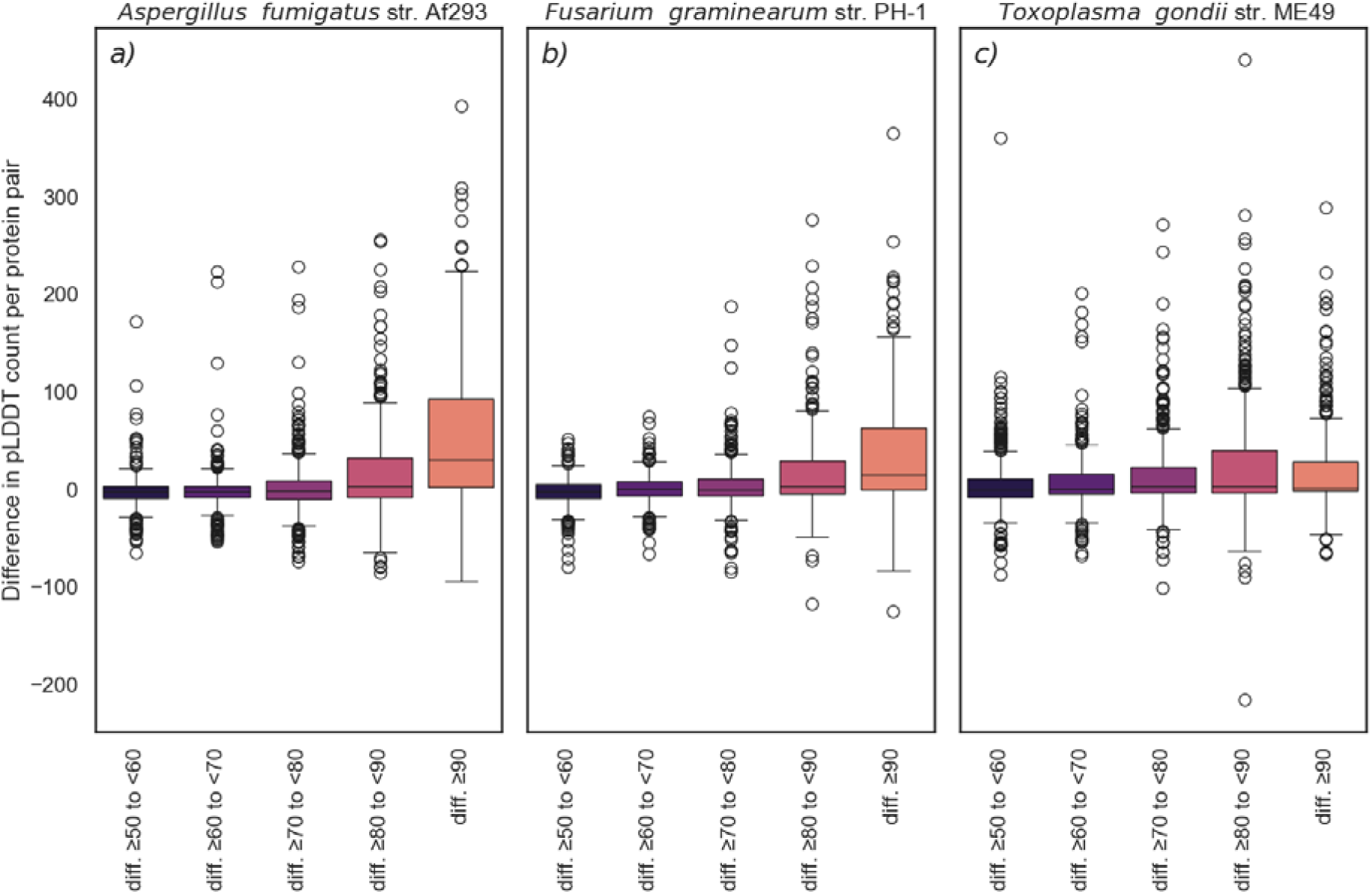
Boxplots of the difference between old and new models in the number of residues that have pLDDT scores within a given range. The box displays the interquartile range (25th percentile, median, 75th percentile), the whiskers represent the 1.5 IQR of the upper and lower quartile and beyond that are outliers.

**Figure 4.**
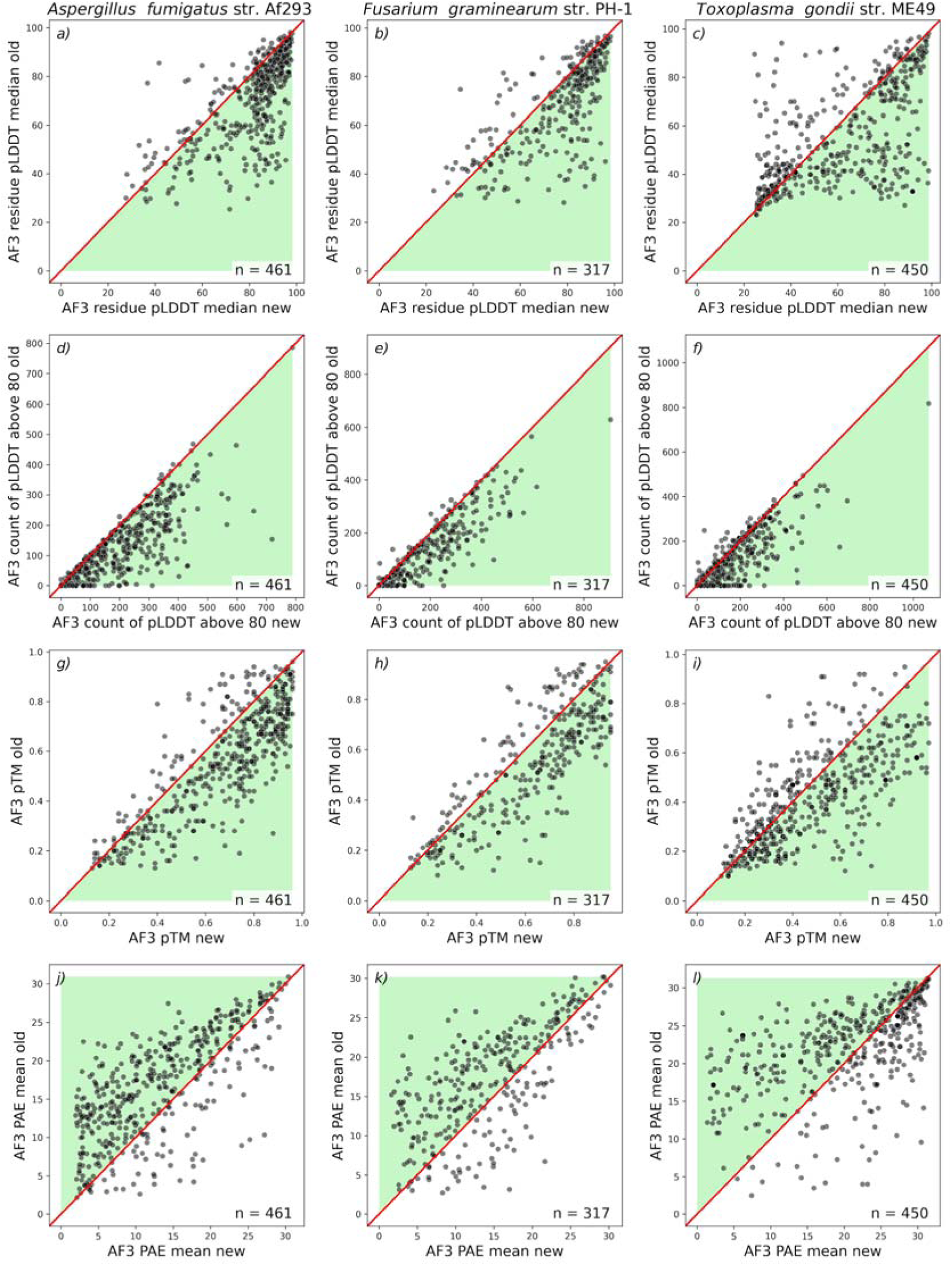
Scatter plots of various AlphaFold 3 scores comparing new models (x axis) to old models (y-axis) The green shading illustrates the direction of positive change from new to old (e.g. an increase in mean pLDDT from old to new is good). The diagonal red line represents no change between old and new models.The number of pairs plotted are displayed in the lower right of each plot.

### Foldseek and InterProScan as supporting tools to AF3

Additional, useful signal may be gleaned from a model’s similarity to experimentally characterised proteins, independently of raw AF3 scores. We used Foldseek (with the PDB database) and InterProScan to provide several additional scores: LDDT, alntmscore, bitscore, and E-value from Foldseek; IPR domain maximum length and IPR domain total length from InterProScan. It should be noted that the number of IPR domains found does increase from old to new models (Supplementary data 1-3), supporting the idea that there is an overall improvement between datasets.

Figure 5 and 6 illustrate a general improvement in most scores, though not in Foldseek’s LDDT or, in a consistent fashion across species, in the alntmscore (Supplementary figure 4). IPR total domain length is the strongest of these additional signals, but it also provides fewer results per gene pair, with IPR domains missing from a third of all gene models (Figure 5 vs 6). For Foldseek, bitscore appears to be the most reliable signal (Figure 2 and 5), and almost all gene models have a hit. However, many of Foldseek’s top hits are not significant, resulting in a similar distribution of results to InterProScan for our two better annotated species: *A. fumigatus* and *F. graminearum* (Figure 2a-b, Supplementary data 1-3). Foldseek contributed best to *T. gondii* which saw larger signals of improvements in its Foldseek scores compared to IPR (Figure 2c).

**Figure 5.**
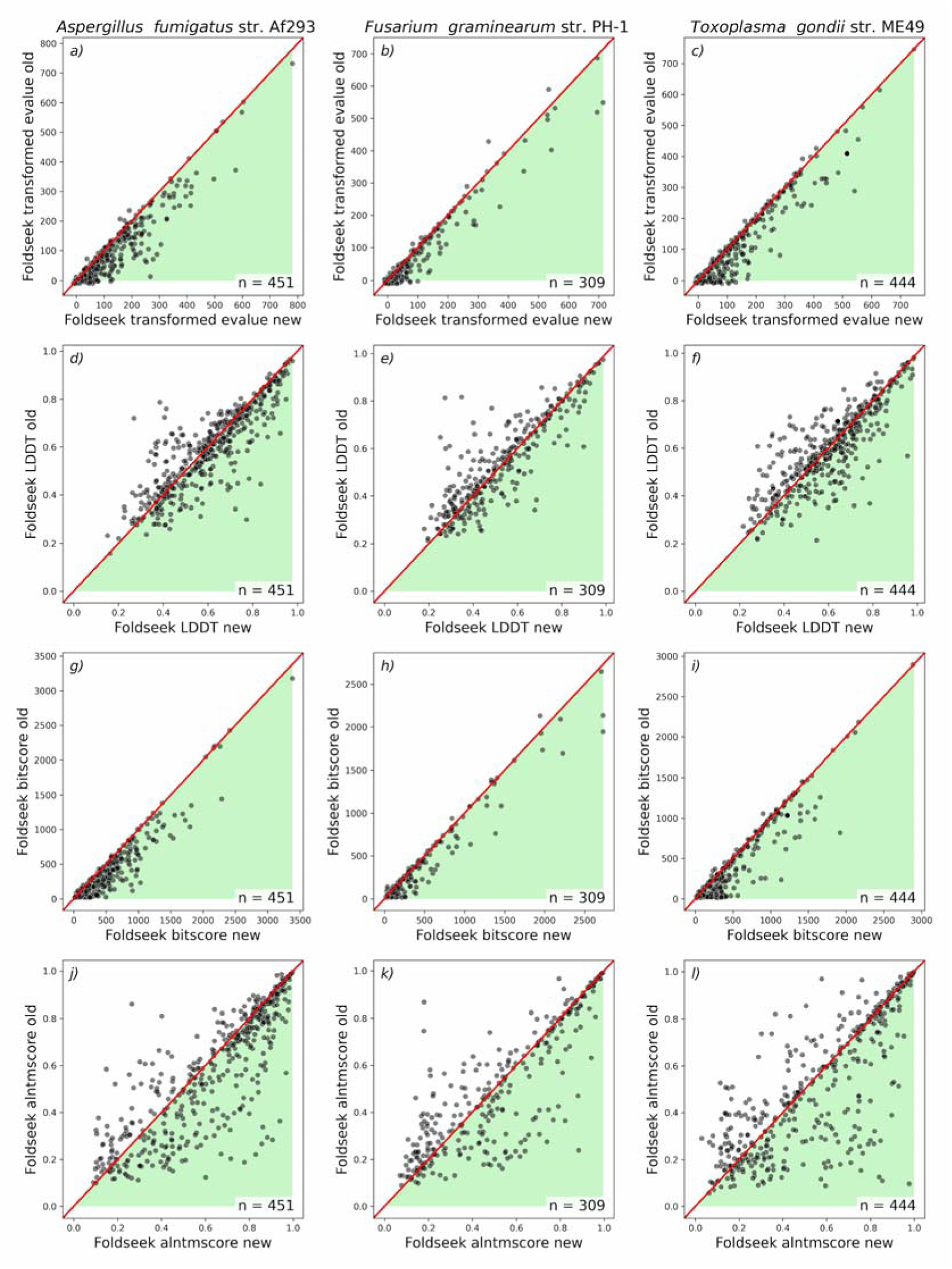
Scatter plots of Foldseek scores comparing new models (x axis) to old models (y-axis) The green shading illustrates the direction of positive change from new to old (e.g. an increase in mean LDDT from old to new is good). The diagonal red line represents no change between old and new models.The number of pairs plotted are displayed in the lower right of each plot.

**Figure 6.**
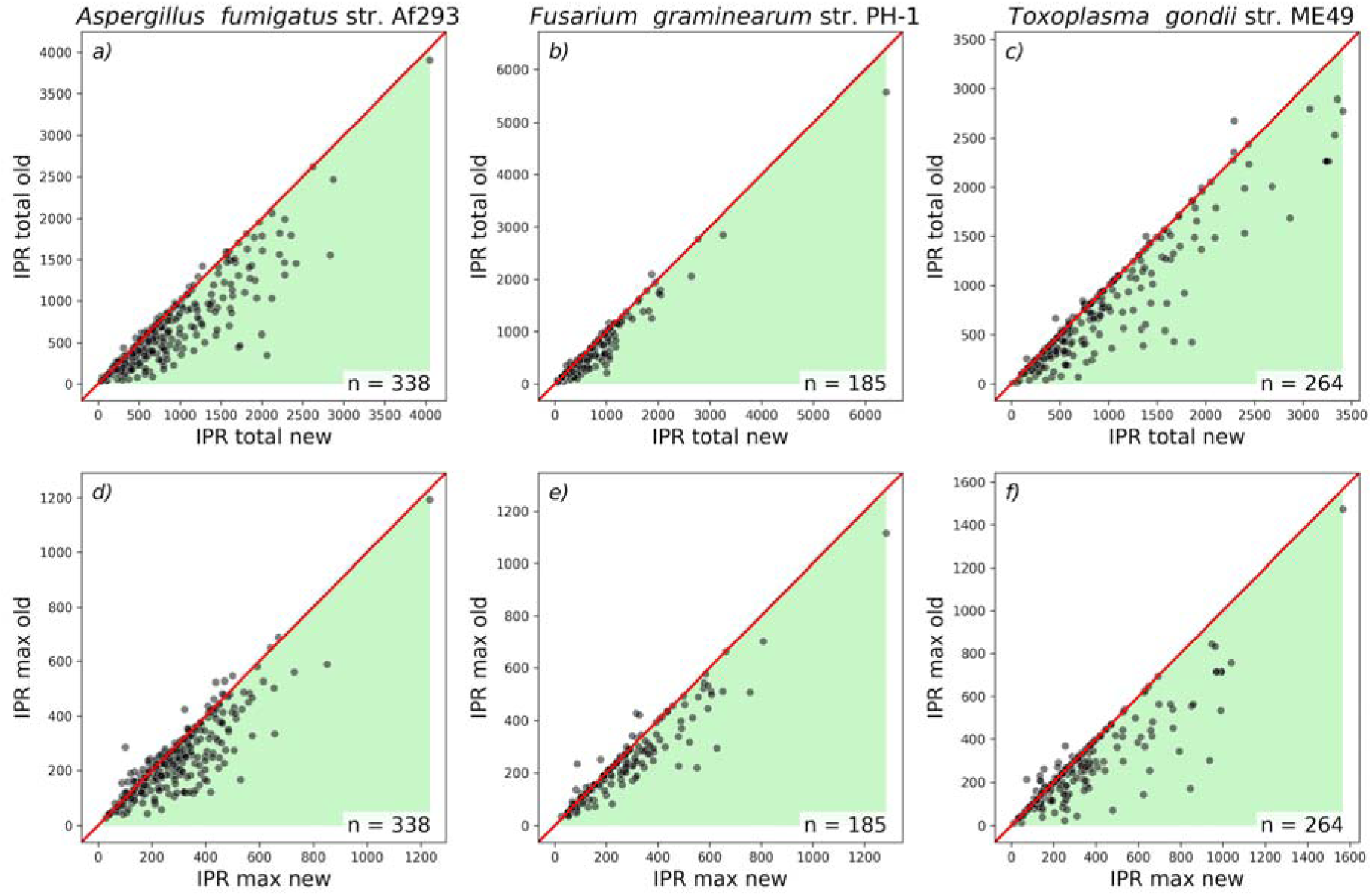
Scatter plots of InterProScan scores comparing new models (x axis) to old models (y-axis) The green shading illustrates the direction of positive change from new to old (e.g. an increase in IPR the maximum domain length score from old to new is good). The diagonal red line represents no change between old and new models. The number of pairs plotted are displayed in the lower right of each plot.

### Combining scores

We next wished to assess whether the AF3, Foldseek and InterProScan scores are redundant, or whether there is a degree of complementarity between them that would encourage integration of all three into future methods. Correlation plots and heat maps illustrate that where AF3 fails to find signal, others do (Supplementary figures 5-7).

Taken in combination, we are able to show a general positive trend in the difference in scores between old and new genes (Figure 2, Supplementary figure 7). Figure 2 shows that combining scores from AF3, Foldseek, and IPR leads to a greater sensitivity to overall model improvement compared to relying on single scores.

### Illustrative cases of success and failure

The bare numbers in Figures 2-6 indicate the value of the scores introduced here, but we additionally illustrate typical successes in Figure 7a and 7b. In these examples, the ‘old’ sequences were derived from misannotated genes containing errors such as missing exons, frameshifts, or incorrect start sites. The corrected annotations in the ‘new’ models yield accurate protein sequences, restoring structural integrity and completing domain matches.

**Figure 7.**
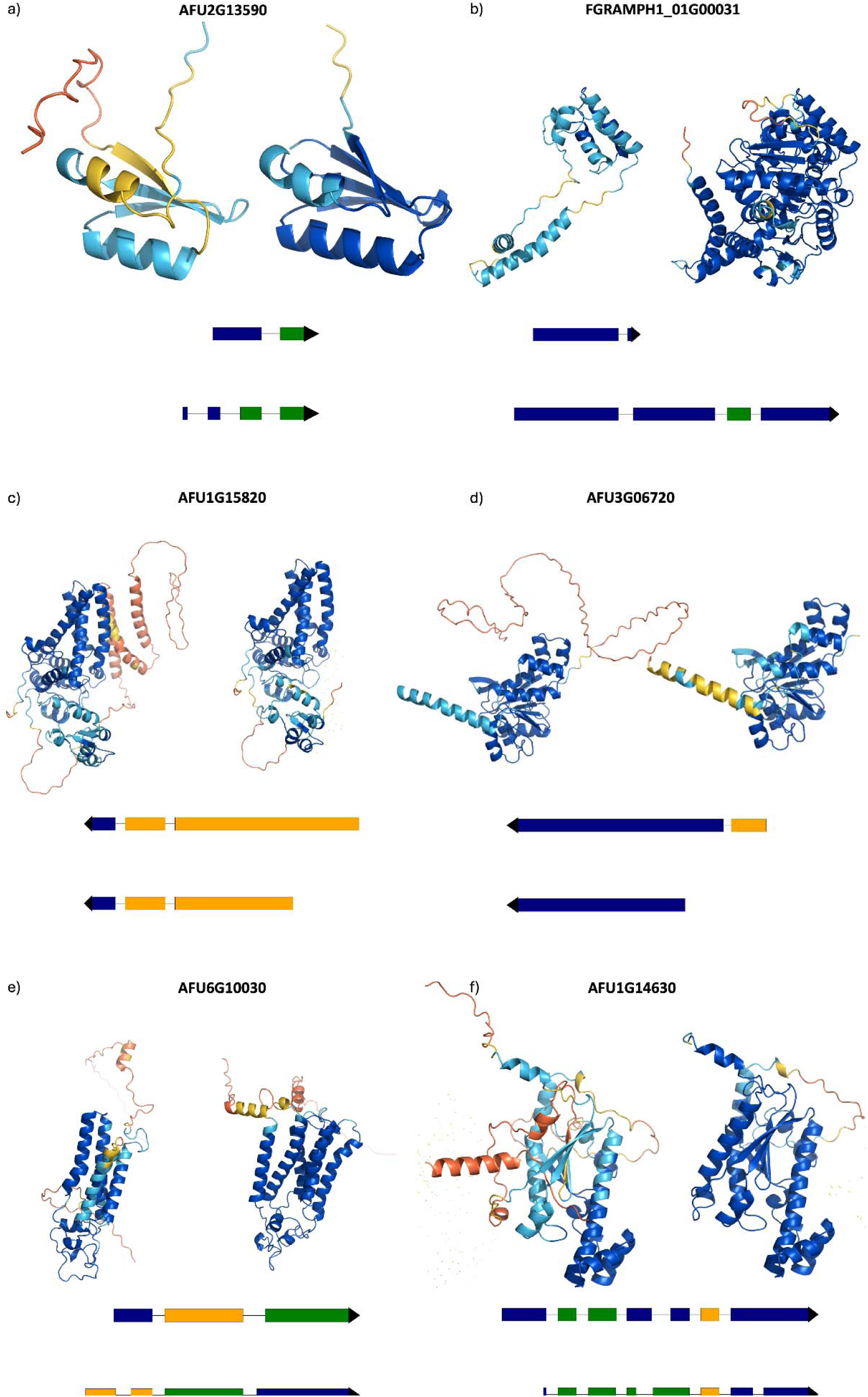
Examples of output protein models for ‘old’ and ‘new’ gene annotations. Each panel shows paired protein models: the ‘old’ model (left) and the corresponding ‘new’ model (right), colored by AlphaFold’s confidence score (pLDDT) from blue (high) to orange (low). Below each pair are schematic representations of the corresponding gene models: the ‘old’ version (top) and the ‘new’ version (bottom). Only coding sequences (CDS) are shown; untranslated regions (UTRs) are omitted for clarity. CDS blocks are coloured according to their reading frame (navy = frame 0, orange = frame 1, green = frame 2), calculated from the GFF phase and lengths of preceding CDS blocks. The black line connects CDS blocks along the genomic locus, with arrowheads indicating the direction of transcription. Differences in the order of CDS colours between the two models indicate potential frameshifts caused by misannotation. The black line represents the genomic locus, with exon schematics aligned to depict their relative positions. Panels (A) and (B) illustrate cases where the ‘new’ models demonstrate improved pLDDT, InterProScan, and Foldseek scores. Panels (C)–(F) show a minority of cases where one or more scores decrease. Cases are discussed in the main text

Interestingly, the example in Figure 7a includes a structural element that is in common between the old and new models, but which corresponds to different sequences in the two cases. Structural alignment of the two models reveals that they are sequence-identical only from residue 20 of the new model, corresponding to residue 31 of the old model. The distinct sequence prior to this region - resulting from the original model being incorrectly predicted as a two exon gene, as opposed to the new model, where it is predicted as a four exon gene - folds to include a superimposable β-strand. The explanation seems to be that AF3, recognising the expected fold from the rest of the sequence, understands that a β-strand should be present and forms it from the most suitable available sequence. However, not being the correct sequence, the strand is recognised as of low-quality and receives low pLDDT scores. Notably, the neighbouring helix in the structure, which *is* composed of the same sequence, is also low-confidence in the old model since the β-strand mentioned above fails to provide a favourable environment for it. Thus, this example illustrates a magnifying effect by which the number of high-confidence residues in the model of an incorrect gene model can be reduced both in the primary region – a translated intron, a frame-shifted translation - but also in neighbouring parts of the structure, enhancing the signal that favours the correct sequence.

In general, the AF3, Foldseek and IPR scores correlate well (Supplementary figure 5) but Figure 7 also illustrates cases of discordance between the scores and what can be learned from them. In Figure 7c, the number of high-confidence residues slightly decreases (from 299 to 288), but the core structure remains accurate, with higher Foldseek and InterProScan scores reflecting better correspondence to known folds and domains, possibly due to more canonical sequence coverage. In Figure 7d, all three metrics decrease slightly (high-confidence residues drop from 229 to 200; Foldseek from 136 to 134; InterProScan from 817 to 762). The difference between old and new annotations is simply an erroneous extra stretch of 106 residues at the N-terminus of the former. Here, the unexpected lower confidence for the helix that starts the correct protein sequence may result from differences, induced by the flanking sequence, in the Multiple Sequence Alignment (MSA) generated for the two cases: it is known that MSA depth is reflected in pLLDT values output by AlphaFold (34).

In Figure 7e, both high-confidence residue count and InterProScan score increase, while Foldseek decreases slightly (from 41 down to 34). The proteins matched by FoldSeek are similar so it appears that the missing helix in the old structure leads to a distortion away from the consensus helical packing of the family. Finally, in Figure 7f, the number of high-confidence residues and the Foldseek score both increase, while the InterProScan score decreases slightly (1173 down to 1113). Despite being manually curated and biologically complete, the ‘new’ sequence may show slightly shifted alignments or small insertions/deletions (e.g., loops or linkers) relative to the statistical domain models used by InterProScan. These subtle differences can lower sequence-based alignment scores even when the overall structure is more accurate.

One important conclusion from examination of these cases is that discordant scores typically change in the wrong direction by only a small amount, suggesting that consideration of the magnitude of the change in a score, not just its direction, should be given more attention in the future. This reflects the fact that all the scores here will be susceptible to some noise.

## Discussion

Manual curation by experts actively improves gene structure models, but it is well known to be a time consuming and expensive task (3,35). Here we explore the idea that scores from 3D structural prediction software could be a useful tool in improving automated gene model quality scoring. These scores could ultimately be used in improving gene annotation quality for many new genomes that are sequenced each year that will likely never see manual curation (36).

Overall, while protein structure-based scores are clearly powerful, our results support the established wisdom that there is no single method of determining the quality of all gene or protein models (37). We chose to focus on AlphaFold 3 and combine its scores with Foldseek (vs the PDB) and InterProScan results to complement pure model-based scores with those derived from a comparison to known protein structural domain data. AlphaFold 3 scores largely support the manual changes made by our curator, and these are further enhanced when combined with other structural information from both Foldseek and InterProScan. InterProScan is powerful on its own, but is limited in scope by the fact that only two thirds of the gene models have a significant hit with an interpro domain. Similarly, Foldseek was only informative for folds currently observed in the PDB. Future work could explore extending the search to TED (The Encyclopedia of Domains; (38)) which captures and classifies recurring domains in the AlphaFold Protein Structure Database (39). The risk of circular logic, by which a protein could be matched to a model of a protein derived from an incorrect gene model, could be minimised by using the newly developed q-score which favours domains that are more ordered and which are seen repeatedly across the database.

One confounding issue identified early on was the impact of intrinsic disorder. Scores like pTM correlate with disorder (Supplementary figure 3), rendering our analyses less likely to define a preferred gene model where differences between old and new gene models fall in disordered regions (although interaction motifs can manifest as islands of higher pLDDT in disordered regions (40)). Among future avenues to explore for disordered regions are the use of a protein language model, as employed by the NetStart 2.0 package for translation start prediction (41). Other approaches could exploit the observation that disordered regions can be annotated with different ‘flavours’, each with distinct functional correlations (42). The simple observation that a protein region has a strong ‘flavour’ may well be a significant sign that it is genuinely translated sequence; but it may also be possible to assess the congruence of the functions of the disorder flavour and the structural domains found in the same protein.

Nevertheless, despite challenges presented for some particular proteins (such as intrinsically disordered), we have demonstrated strong signals and complementary signals from protein structure tools, which can be adopted to aid future software development. The attraction of tools that work directly from a protein fasta file is that these can also help with gene model scoring, in the absence of deep experimental data (e.g. transcriptomics). We envisage that protein structure-derived metrics can be added to improve gene finding pipelines, or post-processing tools that select the best gene model from a panel of alternatives.

## Conclusions

In conclusion, scores derived from the latest structural prediction tools can be highly effective for selection of gene models, but have room for improvement. Combining the outputs of structural prediction tools and gene accuracy measures may be useful in producing quality scores for gene models in general. It will require concerted effort to curate gene model databases for deep learning models that capture the width and breadth of variation in nature, and databases like VEuPathDB that integrate multiomics data are well placed to support this work.

## Methods

### Selection of genomes and gene model pairs

Gene model annotations for *Toxoplasma gondii* str. ME49 underwent substantial changes on ToxoDB (A VEuPathDB sub-site; (43)) between build version 65 and 66. Over 1200 genes were manually curated to improve or correct their structure. Metadata for altered gene models were retrieved from https://toxodb.org/toxo/app/static-content/ToxoDB/news.html#ToxoDB66Released. *Fusarium graminearum* and *Aspergillus fumigatus* underwent similar curation of 2047 and 1969 gene models respectively between build version 68 and a future release of their genomes on FungiDB (44,45). The records of these changes are currently publicly available on the Apollo community annotations track and can be viewed through Jbrowse.

Methods for carrying out annotation can be found in the supplementary information. Files for all of the genomes used in this paper, including protein sequence files, gffs, and metadata, have been included as supplementary information. All data was retrieved from ToxoDB and FungiDB data download pages for relevant build version numbers (65 and 66 for *Toxoplasma gondii*, and 68 and internal data for the two fungi). All updates to gene models were carried out by Böhme through Apollo (46) using transcriptomics data and community annotations publicly available on VEuPathDB.

Once downloaded, protein sequences for new and old annotation were aligned and scored with BioPython version 1.85 (47). Sequences were filtered for pairs that shared <80% amino acid sequence identity and underwent edits classed as ‘changed’ (where boundaries of exons, introns, or UTRs had been amended). This resulted in 444, 317 and 461 gene pairs respectively from *T. gondii* str. ME49*, F. graminearum* str. PH-1 and *A. fumigatus* str. Af293 for new and old versions.

### Protein-based analyses

The sets of gene pairs with <80% sequence identity for each of the three organisms were run through structural prediction software AlphaFold 3 (AF3) (21). Results from AlphaFold 3 were then taken forward and run through Foldseek (30). Both AlphaFold 3 and Foldseek were run on a HPC cluster system on GPU and CPU respectively through singularity containers. Per residue and per protein pLDDT scores were extracted and calculated from the AlphaFold 3 model output in Python.

InterProScan version 5.73-104.0 (28) was run on amino acid coding sequence with all applications, and the flags: --iprlookup --goterms. The output for each gene model was filtered for hits with IPR domains, and from this we took the length of the longest IPR domain (max IPR) and the total length of all IPR domains (total IPR).

Metapredict3 (48) was run on CPU with default options for disorder prediction. Mean disorder was taken for each protein, across individual disorder scores per amino acid.

All other analyses, processing and visualisations were carried out in Python version 3.11.

Prior to plotting or scoring differences between old and new models, the data from Foldseek was masked based on whether the Foldseek hit had a significant E-value below 0.05. When this threshold was exceeded, the E-value was set to 10 (0 signifies an exact match) and all other Foldseek scores were set to 0. The 0.05 threshold was decided upon by analysing the effect of increasing threshold stringency (Supplementary figure 4).

For differences in scores between old and new annotations, where an increase in a score is seen as indicative of an improved gene model, a positive delta (diff) metric will score +1, and negative −1. Where a decrease is seen as indicative of an improved gene model, a positive delta (diff) metric will score −1, and negative +1. The MaxAbs scaling and transformation method from scikit-learn version 1.7.1 (49) was also applied to difference values between old and new models across multiple data sources for analysis and visualisation.

For sums of scores, scores were always summed after applying MaxAbs transformation. Separate transformations were performed for all 11 variables, AF3+Foldseek+IPR, AF3+Foldseek, and AF3+IPR. The best single variables from each program (*’AF3 total residue pLDDT over 80 diff.’, ‘Foldseek bitscore diff.’, ‘IPR domain total length diff.’*) were used to calculate sums for AF3+Foldseek+IPR, AF3+Foldseek, and AF3+IPR. Summed scores were then masked with the same win/lose criteria as previous, where anything >0 was a win (+1) and <0 was a lose (−1).

Code for reproducing the analyses here are in github: https://doi.org/10.5281/zenodo.17379945.

## Abbreviations

AF2: AlphaFold 2
AF3: AlphaFold 3
HPC: High-performance computing
IPR: InterProScan

## Declarations

### Ethics approval and consent to participate

Not applicable

### Consent for publication

Not applicable

### Availability of data and materials

Data used in this publication can be found at FungiDB.

The datasets generated or analysed in this study, along with the code used, can be found on Zenodo with the following dois: 10.5281/zenodo.17287464, 10.5281/zenodo.17290574, 10.5281/zenodo.17290584

### Authors’ contributions

HRD carried out the majority of data processing and analysis, including running Foldseek, Metapredict3, and InterProScan.

HRD, SM, DR, and AJ were major contributors to writing the manuscript, project conceptualisation, and direction.

DR, SM, and AJ contributed to data analysis.

SM generated preliminary data.

PW built containers for and ran AlphaFold 3, and was indispensable in IT expertise.

UB carried out manual curation and construction of metadata files for alterations to existing gene models on Apollo.

## Acknowledgements

Not applicable

## References

1. Mudge JM, Carbonell-Sala S, Diekhans M, Martinez JG, Hunt T, Jungreis I, et al. GENCODE 2025: reference gene annotation for human and mouse. Nucleic Acids Res. 2024 Nov 20;53(D1):D966–75.

2. Salzberg SL. Next-generation genome annotation: we still struggle to get it right. Genome Biol. 2019 May 16;20(1):92.

3. Rhie A, McCarthy SA, Fedrigo O, Damas J, Formenti G, Koren S, et al. Towards complete and error-free genome assemblies of all vertebrate species. Nature. 2021 Apr;592(7856):737–46.

4. Lewin HA, Robinson GE, Kress WJ, Baker WJ, Coddington J, Crandall KA, et al. Earth BioGenome Project: Sequencing life for the future of life. Proceedings of the National Academy of Sciences. 2018 Apr 24;115(17):4325–33.

5. The Darwin Tree of Life Project Consortium. Sequence locally, think globally: The Darwin Tree of Life Project. Proceedings of the National Academy of Sciences of the United States of America. 2022 Jan 18;119(4):e2115642118.

6. Lukashin AV, Borodovsky M. GeneMark.hmm: New solutions for gene finding. Nucleic Acids Res. 1998 Feb 1;26(4):1107–15.

7. Brůna T, Lomsadze A, Borodovsky M. GeneMark-ETP significantly improves the accuracy of automatic annotation of large eukaryotic genomes. Genome Res. 2024 Jun 25;34(5):757–68.

8. Burge C, Karlin S. Prediction of complete gene structures in human genomic DNA. J Mol Biol. 1997 Apr 25;268(1):78–94.

9. Stanke M, Steinkamp R, Waack S, Morgenstern B. AUGUSTUS: a web server for gene finding in eukaryotes. Nucleic Acids Research. 2004 Jul 1;32(Web Server issue):W309.

10. Cantarel BL, Korf I, Robb SMC, Parra G, Ross E, Moore B, et al. MAKER: An easy-to-use annotation pipeline designed for emerging model organism genomes. Genome Res. 2008 Jan 1;18(1):188–96.

11. Gabriel L, Brůna T, Hoff KJ, Ebel M, Lomsadze A, Borodovsky M, et al. BRAKER3: Fully automated genome annotation using RNA-seq and protein evidence with GeneMark-ETP, AUGUSTUS, and TSEBRA. Genome Res. 2024 May 1;34(5):769–77.

12. The NCBI Eukaryotic Genome Annotation Pipeline [Internet]. [cited 2025 Oct 1]. Available from: https://www.ncbi.nlm.nih.gov/refseq/annotation_euk/process/

13. Aken BL, Ayling S, Barrell D, Clarke L, Curwen V, Fairley S, et al. The Ensembl gene annotation system. Database: The Journal of Biological Databases and Curation. 2016 Jun 23;2016:baw093.

14. Stiehler F, Steinborn M, Scholz S, Dey D, Weber APM, Denton AK. Helixer: cross-species gene annotation of large eukaryotic genomes using deep learning. Bioinformatics. 2020 Dec 16;36(22-23):5291–8.

15. Gabriel L, Hoff KJ, Brůna T, Borodovsky M, Stanke M. TSEBRA: transcript selector for BRAKER. BMC Bioinformatics. 2021 Nov 25;22(1):1–12.

16. Libouban R. Comparison of two annotation tools - Helixer and Braker3. 2025 Jun 25 [cited 2025 Oct 15]; Available from: https://github.com/galaxyproject/training-material

17. Banerjee S, Bhandary P, Woodhouse M, Sen TZ, Wise RP, Andorf CM. FINDER: an automated software package to annotate eukaryotic genes from RNA-Seq data and associated protein sequences. BMC Bioinformatics. 2021 Apr 20;22(1):205.

18. Holt C, Yandell M. MAKER2: an annotation pipeline and genome-database management tool for second-generation genome projects. BMC Bioinformatics. 2011 Dec 22;12:491.

19. Olson AJ, Ware D. Ranked choice voting for representative transcripts with TRaCE. Bioinformatics. 2021 Jul 23;38(1):261–4.

20. Sommer MJ, Zimin AV, Salzberg SL. PSAURON: a tool for assessing protein annotation across a broad range of species. NAR Genom Bioinform. 2025 Mar;7(1):lqae189.

21. Abramson J, Adler J, Dunger J, Evans R, Green T, Pritzel A, et al. Accurate structure prediction of biomolecular interactions with AlphaFold 3. Nature. 2024 May 8;630(8016):493–500.

22. Jumper J, Evans R, Pritzel A, Green T, Figurnov M, Ronneberger O, et al. Highly accurate protein structure prediction with AlphaFold. Nature. 2021 Jul 15;596(7873):583–9.

23. Abbass J. From CASP13 to the Nobel Prize: DeepMind’s AlphaFold Journey in Revolutionizing Protein Structure Prediction and Beyond. Curr Protein Pept Sci [Internet]. 2025 Oct 2; Available from: 10.2174/0113892037374986250711152300

24. Ruff KM, Pappu RV. AlphaFold and Implications for Intrinsically Disordered Proteins. J Mol Biol. 2021 Oct 1;433(20):167208.

25. Dunbrack RL Jr. : What’s wrong with AlphaFold’s score and how to fix it [Internet]. bioRxiv. 2025. Available from: 10.1101/2025.02.10.637595

26. Lomize MA, Pogozheva ID, Joo H, Mosberg HI, Lomize AL. OPM database and PPM web server: resources for positioning of proteins in membranes. Nucleic Acids Res. 2012 Jan;40(Database issue):D370–6.

27. Berman H, Henrick K, Nakamura H. Announcing the worldwide Protein Data Bank. Nat Struct Biol. 2003 Dec;10(12):980.

28. Jones P, Binns D, Chang HY, Fraser M, Li W, McAnulla C, et al. InterProScan 5: genome-scale protein function classification. Bioinformatics. 2014 May 1;30(9):1236–40.

29. Blum M, Andreeva A, Florentino LC, Chuguransky SR, Grego T, Hobbs E, et al. InterPro: the protein sequence classification resource in 2025. Nucleic Acids Res. 2025 Jan 6;53(D1):D444–56.

30. van Kempen M, Kim SS, Tumescheit C, Mirdita M, Lee J, Gilchrist CLM, et al. Fast and accurate protein structure search with Foldseek. Nature Biotechnology. 2023 May 8;42(2):243–6.

31. Alvarez-Jarreta J, Amos B, Aurrecoechea C, Bah S, Barba M, Barreto A, et al. VEuPathDB: the eukaryotic pathogen, vector and host bioinformatics resource center in 2023. Nucleic Acids Res. 2024 Jan 5;52(D1):D808–16.

32. Feng ZP, Zhang X, Han P, Arora N, Anders RF, Norton RS. Abundance of intrinsically unstructured proteins in P. falciparum and other apicomplexan parasite proteomes. Mol Biochem Parasitol. 2006 Dec;150(2):256–67.

33. Zaccaron AZ, Stergiopoulos I. The dynamics of fungal genome organization and its impact on host adaptation and antifungal resistance. J Genet Genomics. 2025 May;52(5):628–40.

34. Costa F, Blum M, Bateman A. Keeping it in the family: using protein family templates to rescue low confidence AlphaFold2 models. Bioinform Adv. 2024 Nov 25;4(1):vbae188.

35. Miga KH, Koren S, Rhie A, Vollger MR, Gershman A, Bzikadze A, et al. Telomere-to-telomere assembly of a complete human X chromosome. Nature. 2020 Sep;585(7823):79–84.

36. GenBank and WGS Statistics [Internet]. [cited 2025 Oct 15]. Available from: https://www.ncbi.nlm.nih.gov/genbank/statistics/

37. Olechnovič K, Monastyrskyy B, Kryshtafovych A, Venclovas Č. Comparative analysis of methods for evaluation of protein models against native structures. Bioinformatics. 2019 Mar 15;35(6):937–44.

38. Lau AM, Bordin N, Kandathil SM, Sillitoe I, Waman VP, Wells J, et al. Exploring structural diversity across the protein universe with The Encyclopedia of Domains. Science. 2024 Nov;386(6721):eadq4946.

39. Varadi M, Bertoni D, Magana P, Paramval U, Pidruchna I, Radhakrishnan M, et al. AlphaFold Protein Structure Database in 2024: providing structure coverage for over 214 million protein sequences. Nucleic Acids Res. 2024 Jan 5;52(D1):D368–75.

40. Piovesan D, Del Conte A, Clementel D, Monzon AM, Bevilacqua M, Aspromonte MC, et al. MobiDB: 10 years of intrinsically disordered proteins. Nucleic Acids Res. 2023 Jan 6;51(D1):D438–44.

41. Nielsen LS, Pedersen AG, Winther O, Nielsen H. NetStart 2.0: prediction of eukaryotic translation initiation sites using a protein language model. BMC Bioinformatics. 2025 Aug 19;26(1):216.

42. Vucetic S, Brown CJ, Dunker AK, Obradovic Z. Flavors of protein disorder. Proteins. 2003 Sep 1;52(4):573–84.

43. Gajria B, Bahl A, Brestelli J, Dommer J, Fischer S, Gao X, et al. ToxoDB: an integrated Toxoplasma gondii database resource. Nucleic Acids Res. 2008 Jan;36(Database issue):D553–6.

44. Basenko EY, Shanmugasundram A, Böhme U, Starns D, Wilkinson PA, Davison HR, et al. What is new in FungiDB: a web-based bioinformatics platform for omics-scale data analysis for fungal and oomycete species. Genetics [Internet]. 2024 May 7;227(1). Available from: 10.1093/genetics/iyae035

45. Basenko EY, Pulman JA, Shanmugasundram A, Harb OS, Crouch K, Starns D, et al. FungiDB: An Integrated Bioinformatic Resource for Fungi and Oomycetes. J Fungi (Basel) [Internet]. 2018 Mar 20;4(1). Available from: 10.3390/jof4010039

46. Lee E, Helt GA, Reese JT, Munoz-Torres MC, Childers CP, Buels RM, et al. Web Apollo: a web-based genomic annotation editing platform. Genome Biology. 2013 Aug 30;14(8):1–13.

47. Cock PJA, Antao T, Chang JT, Chapman BA, Cox CJ, Dalke A, et al. Biopython: freely available Python tools for computational molecular biology and bioinformatics. Bioinformatics. 2009 Jun 1;25(11):1422–3.

48. Emenecker RJ, Griffith D, Holehouse AS. Metapredict: a fast, accurate, and easy-to-use predictor of consensus disorder and structure. Biophys J. 2021 Oct 19;120(20):4312–9.

49. Pedregosa F, Varoquaux G, Gramfort A, Michel V, Thirion B, Grisel O, et al. Scikit-learn: Machine Learning in Python. Journal of Machine Learning Research. 2011;12(85):2825–30.

