## Supplementary Material for "The promise of AlphaFold for gene structure annotation"

### Institution addresses

^1^ Department of Biochemistry, Cell and Systems Biology, Institute of Systems, Molecular and Integrative Biology, University of Liverpool, United Kingdom

^2^ Computational Biology Facility, University of Liverpool, United Kingdom

^3^ Department of Biology, University of Pennsylvania, Philadelphia, PA 19104, USA

### Supplementary data

**Supplementary data 1**, *Aspergillus fumigatus* data files (AlphaFold 3, Metapredict3, InterProScan, Foldseek, fasta, gff, and summary of structural changes) - 10.5281/zenodo.17287464

**Supplementary data 2**, *Fusarium graminearum* data files (AlphaFold 3, Metapredict3, InterProScan, Foldseek, fasta, gff, and summary of structural changes) - 10.5281/zenodo.17290574

**Supplementary data 3**, *Toxoplasma gondii* data files data files (AlphaFold 3, Metapredict3, InterProScan, Foldseek, fasta, gff, and summary of structural changes) - 10.5281/zenodo.17290584

### Supplementary Methods

##### Structural Annotation of *Toxoplasma gondii* ME49, *Aspergillus fumigatus* Af293 and *Fusarium graminearum* PH-1.

Manual structural curation of gene models was performed using the Apollo graphical browser-based curation tool (version 2.6.7) (46) integrated within VEuPathDB. Gene models were examined using RNA-Seq evidence tracks available in VEuPathDB to refine structural features, including exon-intron boundaries, start codon positions, and isoform annotations. Additional edits involved merging or splitting gene models and adding new gene models.

Once structural annotation in Apollo was finalised, a patch build process was initiated. The first process that happens is the gene model diff script is initially run (<https://github.com/VEuPathDB/gene_model_diff>) on both, the core database of the original gene set and the dumped apollo annotation file. Only models that are tagged with a finished status are parsed through. The gene model diff provides an initial quality control mechanism that flags problematic annotations (models that are devoid of start and stop codons against the logic of a gene model) and summarises the changes whether models are new, splits, merges, modified or the same. During the patch build process split, merged and new genes will get a new gene/transcript ID.

### Supplementary figures

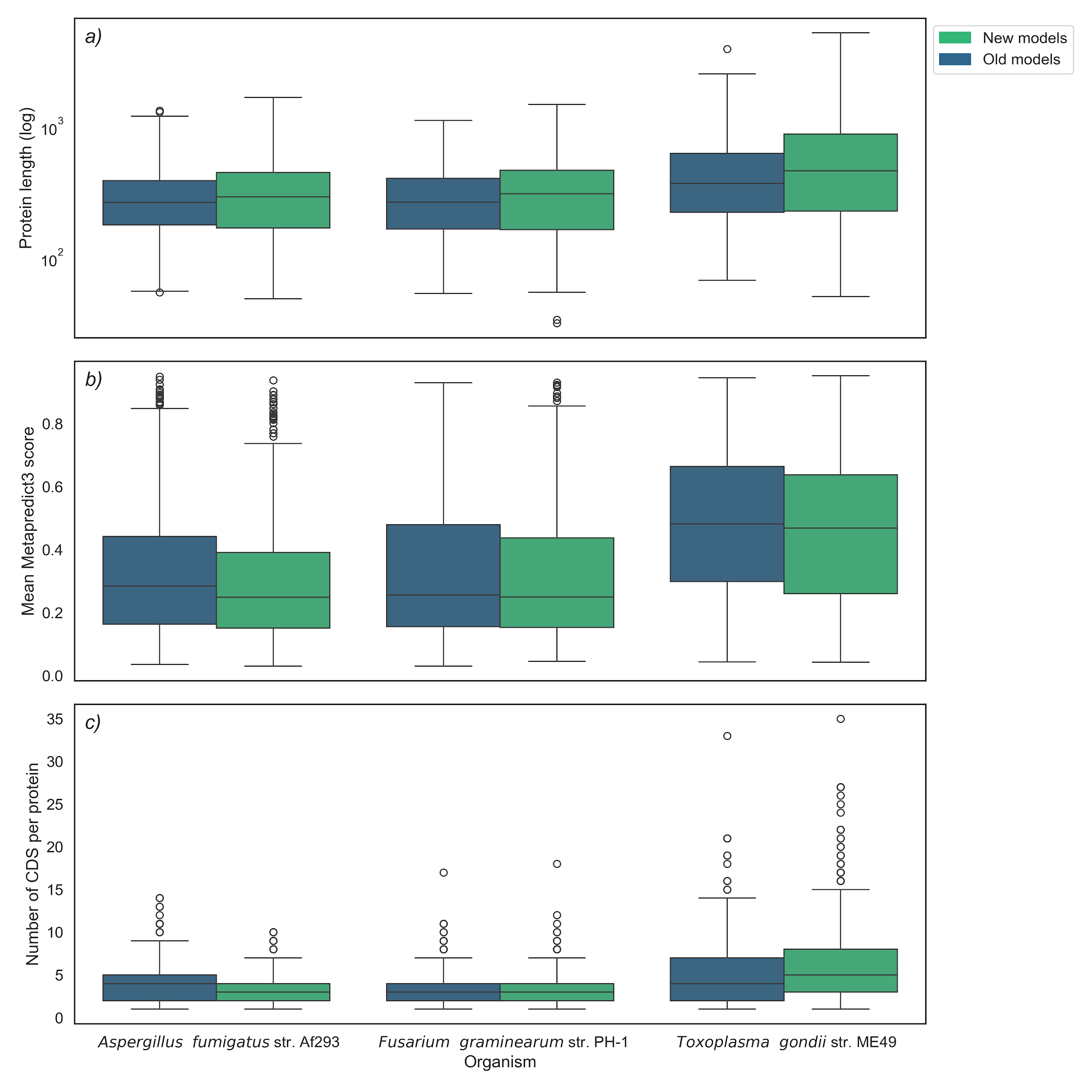

***Supplementary figure 1. Descriptive summary comparison between old and new annotations for all three species,*** *illustrating the change in a) protein length, b) mean Metapredict3 score, and c) the change in the number of coding DNA sequences (CDS) per protein.*

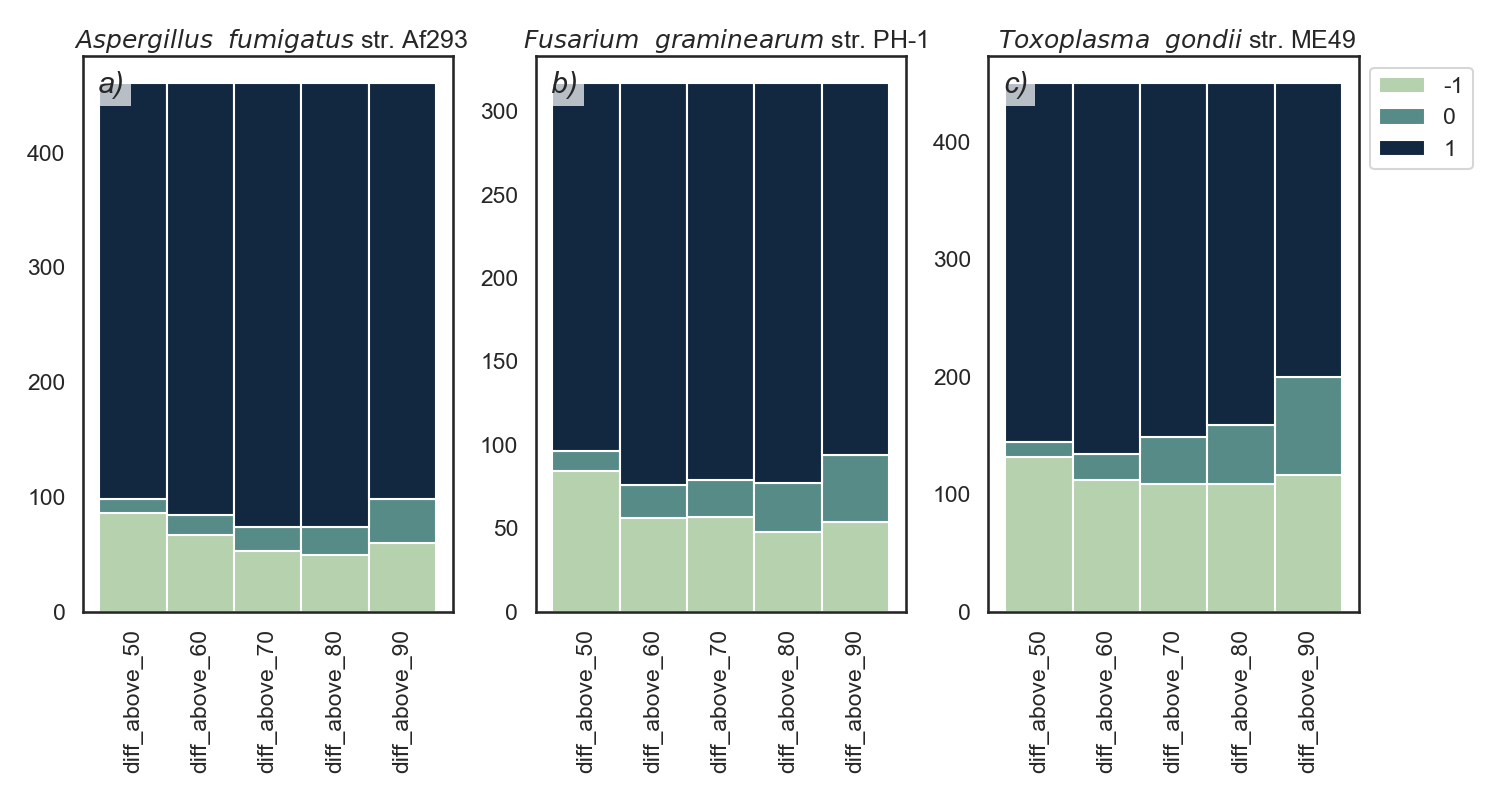

***Supplementary figure 2. win/loss stacked bars for pLDDT counts above a threshold.*** *A win (+1) shows where a new model has more pLDDT residues scoring above the threshold listed on the x axis. A draw (0) is where the number of pLDDT residues scoring above the threshold are the same. A loss (-1) is where the new model has fewer pLDDT residues scoring above the threshold.*

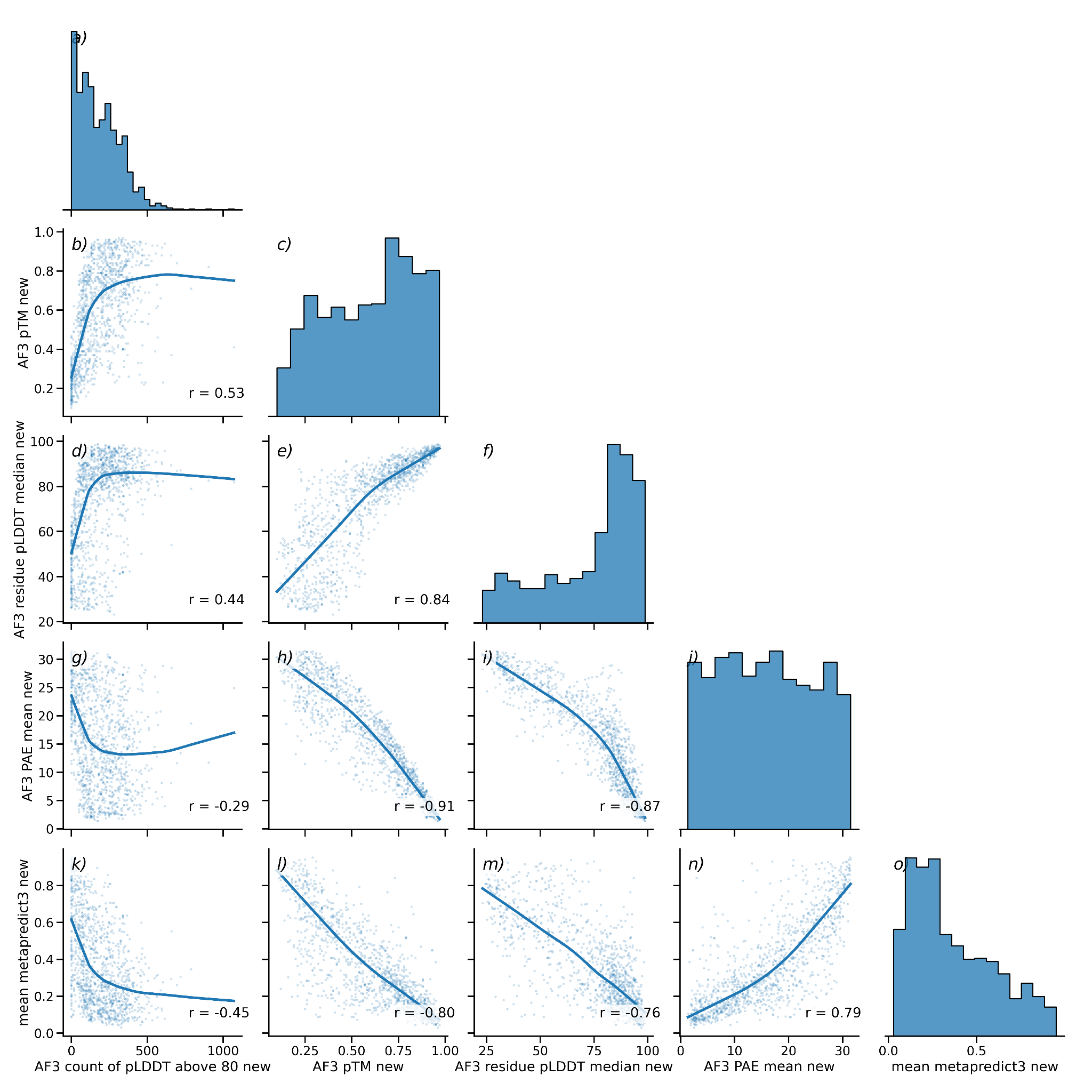

***Supplementary figure 3. Paired scatter plots of mean Metapredict3 disorder scores, AF3, PAE, pTM, pLDDT median, and pLDDT > 80.*** *Scores across all three species for new models are shown. A LOWESS regression line is shown in solid blue, and a Pearson's r score is displayed for each correlation. A histogram for each of the five scores is shown on the diagonal.*

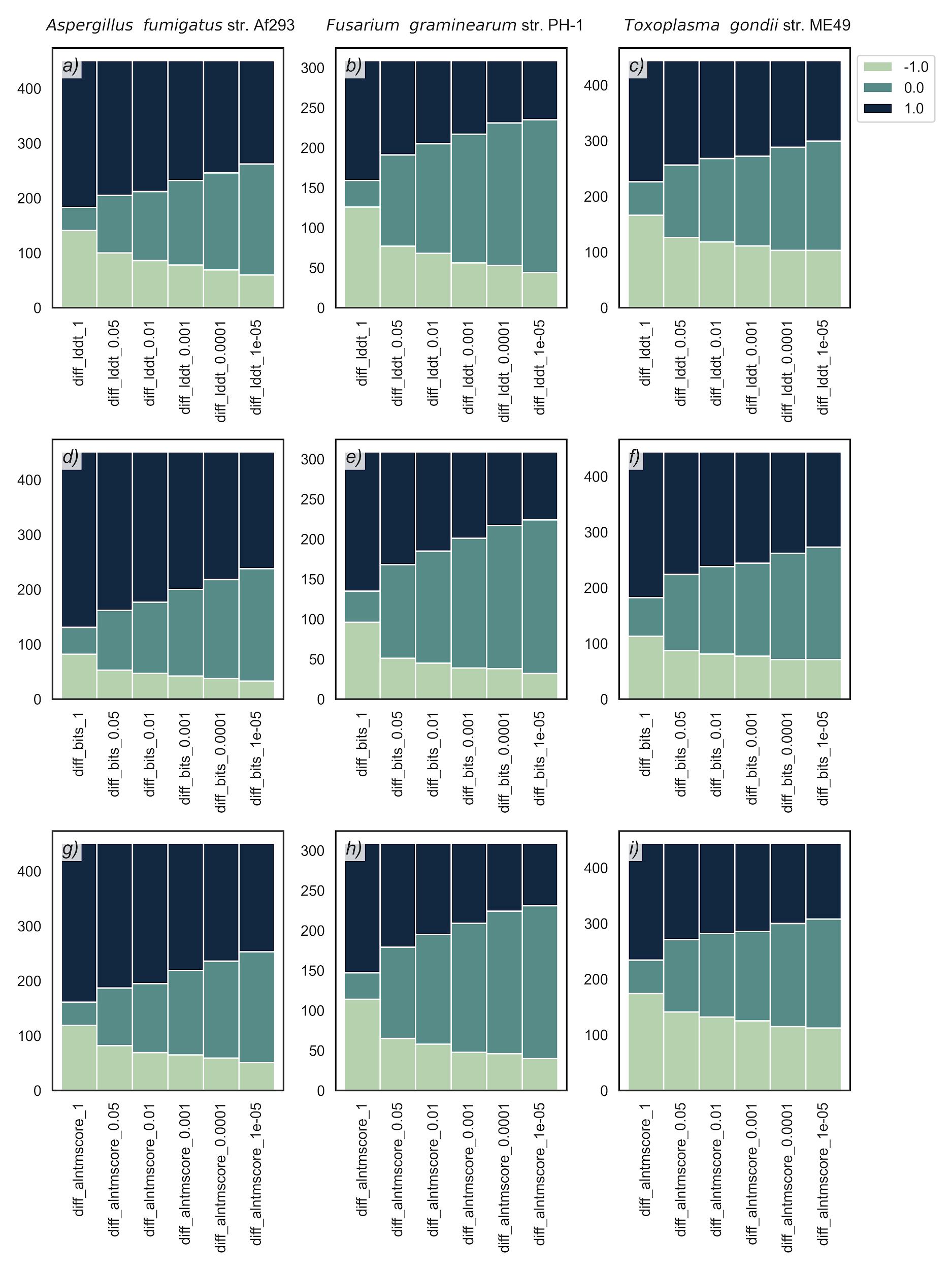

***Supplementary figure 4. win/loss stacked bars for Foldseek scores masked by E-value threshold.*** *Foldseek scores were masked based on whether the E-value was significant. For instance, for an E-value threshold of 0.05, all proteins with E-values greater than 0.05 would be set to zero. Wins and losses were calculated from the differences in these new masked values and scored.* *A win (+1) shows where a new model scores higher than the old model. A draw (0) is where models score the same. A loss (-1) is where the new model scores less than the old.*

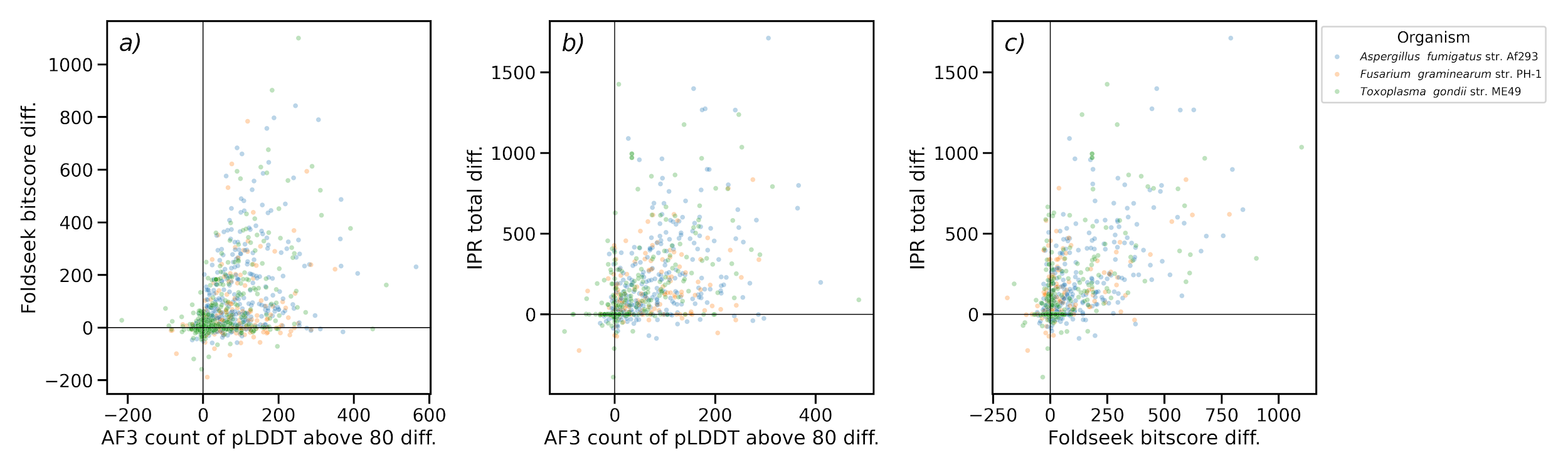

***Supplementary figure 5. The correlation between the three best scores (pLDDT > 80, Bitscore, IPR total length).*** *Scores across all three species for new models are shown. Black lines indicate zero on the x and y axis.*

***Supplementary figure 6a. A clustered heatmap of the MaxAbs transformed difference values for Aspergillus fumigatus str. Af293.*** *In all cases pink represents a positive change from old to new models, blue represents a negative change. Scores where a smaller value is better (e.g. PAE) have been flipped to match this by -1 multiplication .*
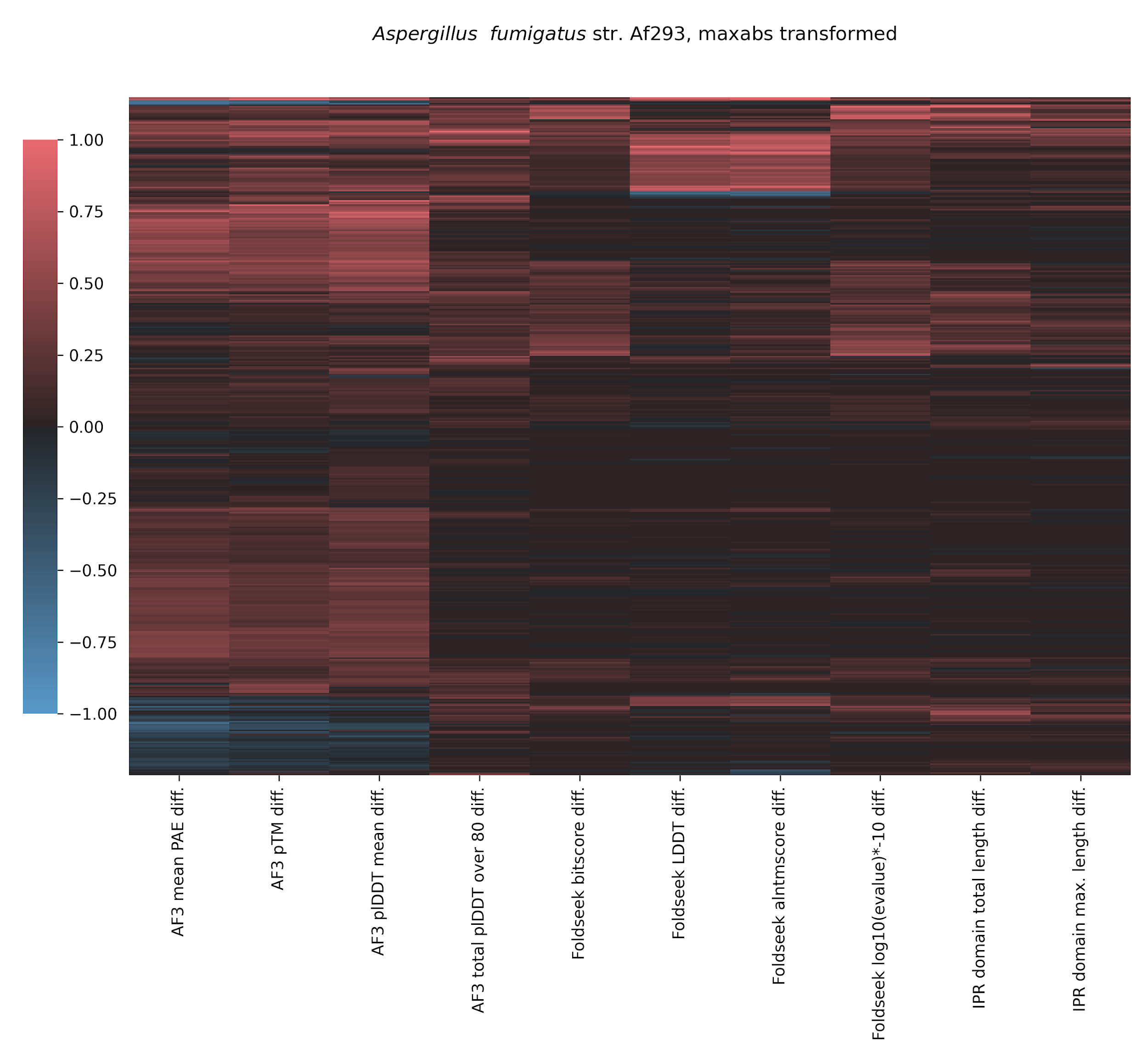

***Supplementary figure 6b. A clustered heatmap of the MaxAbs transformed difference values for Fusarium graminearum str. PH-1.*** *In all cases pink represents a positive change from old to new models, blue represents a negative change. Scores where a smaller value is better (e.g. PAE) have been flipped to match this by -1 multiplication .*
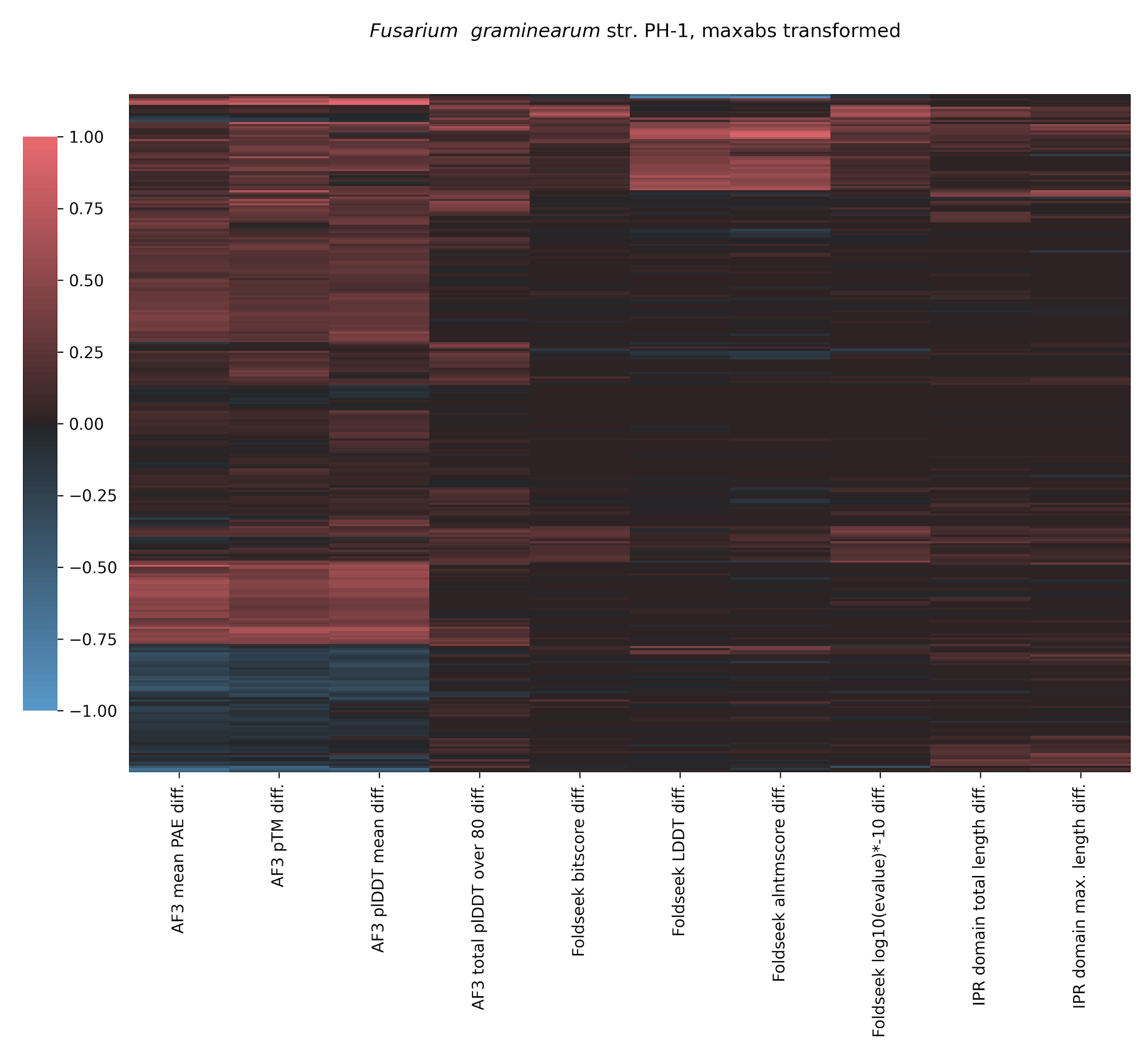

***Supplementary figure 6c. A clustered heatmap of the MaxAbs transformed difference values for Toxoplasma gondii str. ME49.*** *In all cases pink represents a positive change from old to new models, blue represents a negative change. Scores where a smaller value is better (e.g. PAE) have been flipped to match this by -1 multiplication.*
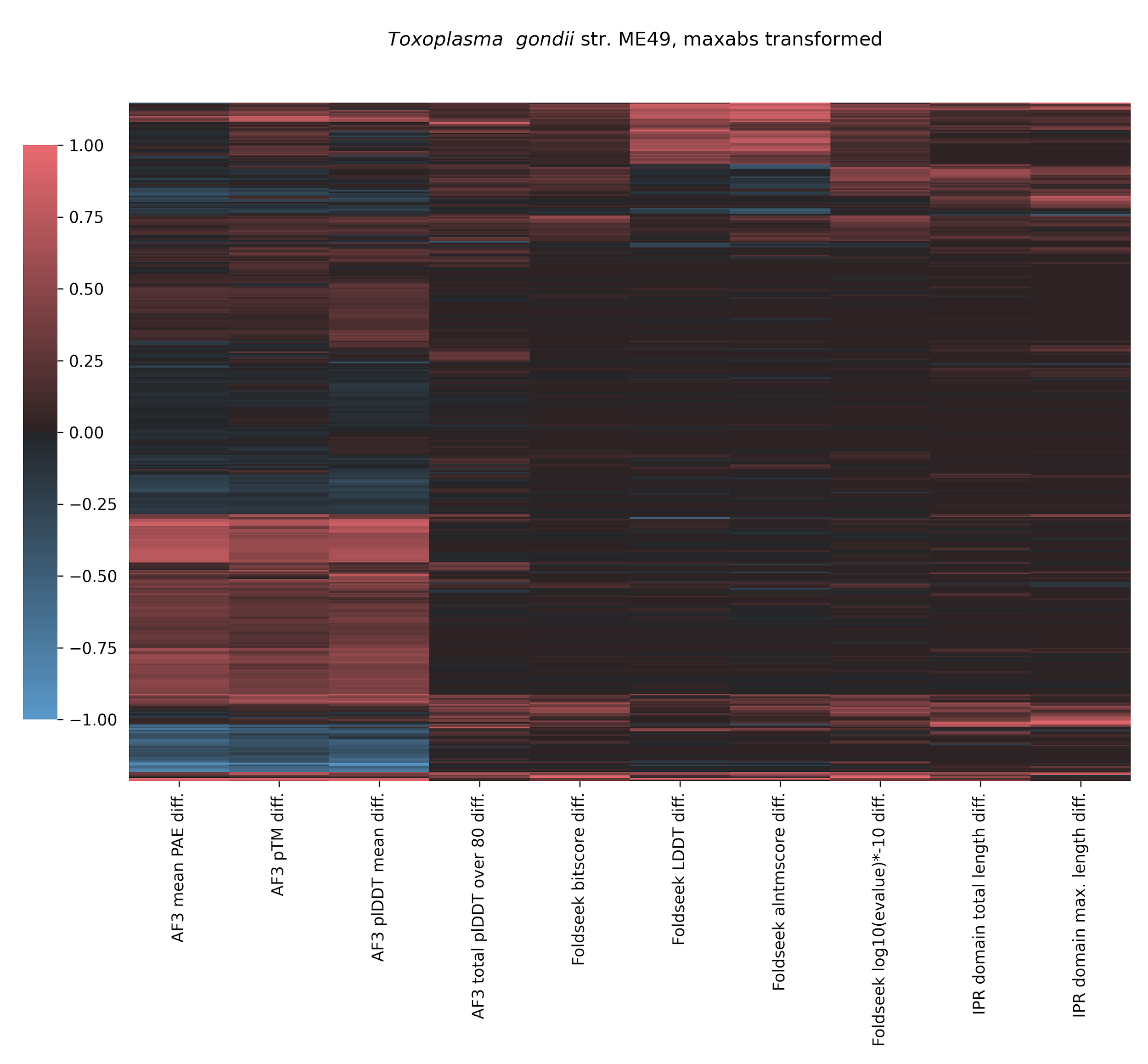

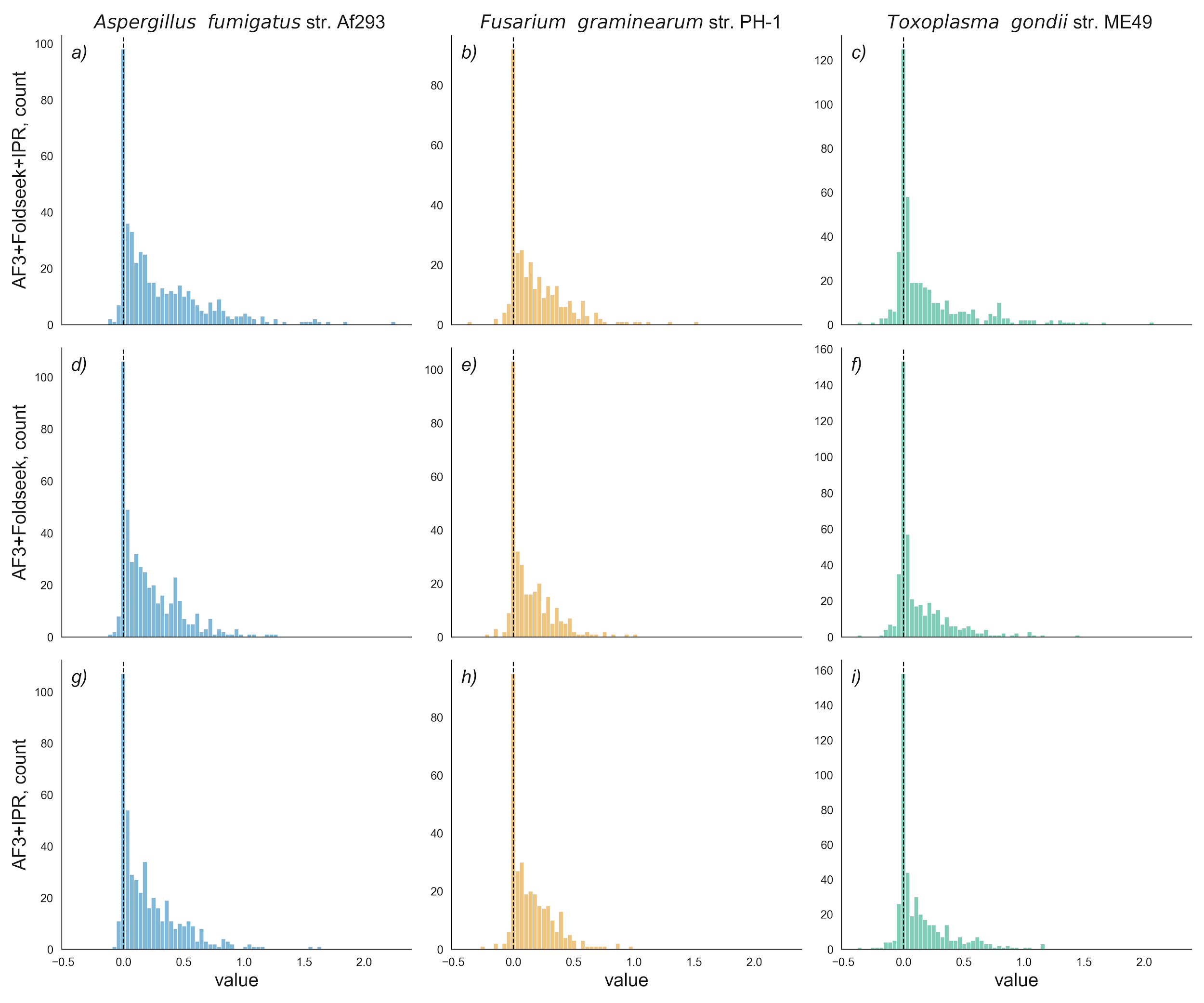

***Supplementary figure 7. Distribution of summed MaxAbs transformed difference scores.*** *Each row displays a different combination of the three best scores (AF3 = pLDDT >80, IPR = IPR total length, Foldseek = Bitscore). Each column displays one species. The black dotted line indicates zero.*
